# Pyntheon: A Functional Analysis Framework for Protein Modifications and Mutations of 83 Model Organisms

**DOI:** 10.1101/725846

**Authors:** Ahmed Arslan

**Affiliations:** Department of Anesthesia, School of Medicine, Stanford University, 300 Pasteur dr, Palo Alto, CA 94305, United States

## Abstract

**Motivation:** Posttranslational modifications (PTMs) modulate proteins activity depending on the dynamics of cellular conditions, in the highly regulated processes that control the reversible nature of these modifications and a cellular state. Due to the unique importance of PTMs, a number of resources are available to analyze the protein modification data for different organisms. These databases are quite informative on a limited number of popular organisms, mostly human and yeast. However there has not been a single database to date that makes it possible to analyze the modified protein residue data for up to 83 model organisms. Moreover, there are limited resources that rely on both protein mutations and modifications in evaluating a phenotype.

**Results:** I am presenting a comprehensive python tool *Pyntheon* that enables users to analyze protein modifications and mutations data. This resource can be used in different ways to know: (i) if the proteins of interest have modifications and (ii) if the modified residues overlap with mutated sites. Additional functions include, analyzing if a PTM-site is present in a functional protein region, like domain and structural regions. In summary, Pyntheon makes it possible for a larger community of researchers to evaluate their curated proteomics data and interpret the impact of mutations on phenotypes.

**Conclusion:** Pyntheon has multifold functions that can help analyzing the protein mutations impact on the modified residues for a large number of popular model organisms.

**Code-Availability:** https://github.com/AhmedArslan/pyntheon

## 1 Introduction

The Post-translational Modifications (PTMs) help proteins to achieve functional trade off with the environment by either activating or inactivating their functions. These modifications are extremely important for the proper protein functions under an ever-fluctuating cellular environment. Any perturbation that leads to either quantitative or qualitative differences in a modification, can result in a phenotypic abnormality (Reimand J. et al. 2105). Several seminal studies have shown that various human diseases have mutated modified residue(s). Besides human, other popular model organisms have greatly instrumented our understanding of protein modifications, both in normal and disease state. Commonly, such studies are conducted either at an individual cell level, like in the case of yeast, or at a whole organism level, such as mice, and have shown the importance of different PTM-types in signaling, transportation, protein attachments. Overall, biological systems studies are a significant step forward in understanding protein-protein interaction and the role of PTMs in these interactions.

The boom in the generation of protein modification data led to the development of several computational tools and curated databases. These tools have been very useful in facilitating in further understanding of the role of PTMs. These resources contain data of limited PTM types (and number of organisms), such as Phosphosite Plus (Hornbeck P. V. et al 2015), PTMcode (Minguez P. et al. 2012) and PTMfunc (Beltrao P. 2012). Despite these resources, there is a lack of tools that contain information for a large number of model organisms (being studied in the lab) that can facilitate scientists in accessing and analyzing the PTMs. Additionally, except for few options (Arslan A and van Noort 2017), most of the available resources do not provide functional assessment of protein mutations impacting the PTMs. Likewise, an automated evaluation tool of mutated PTM residues present in different protein functional regions, like domain or structural features, is lacking.

To fill this gap, I developed Pyntheon, a python package that can help users from across the scientific spectrum in analyzing the PTMs data from 83 popular model organisms (see “species_id.txt” for list of all the available organisms). The command-line tool allows users to (i) retrieve the PTMs data locally for the proteins of 83 different organisms; (ii) analyze if protein residues of interest (mutations) overlap with the PTMs sites; and (iii) if the PTMs of interest are present in a protein functional region like domain or helix.

To develop the database, I retrieved protein modification data from UniProt (The UniProt Consort. 2019) and dbPTM (Lee TY et al 2006) for 83 common model organisms. I compiled the non-redundant data for each species and curated all available PTM information with respect to their presence in different functional regions. A non-redundant PTM database contained 248,066 PTM-sites.

To develop the tool, I implemented this method in python (version 2.x and 3.x) as it makes the tool easy to use: a simple one-line command allows accessing and analyzing the compiled PTMs data (see supplementary information). This type of methods has many benefits over conventional html-based tools. It allows users to retrieve big data as flat-text files and is a much faster way to analyze the user-provided data than the conventional tools.

## 2 Examples

The two following examples show the utility and robustness of Pyntheon. In the first case, I retrieved protein mutation data on human cancers from COSMIC database (Forbes et al 2014). I selected 262,013 non-redundant pathogenic amino-acid alterations present in 7,574 proteins from different tumors (see supplementary information). With a simple one-line command (A) I mapped all the tumor-associated mutational data to human PTMs data and conveniently observed (i) how many proteins that are known in human tumors are modified and (ii) how many of these protein mutation positions overlap with the modified residues.

### $ python3 pyntheon.py -f file.txt -s hs -m TRUE …… (A)

Out of the tumor genes analyzed, 746 proteins have different types of modifications, with phosphorylation being the most abundant. That is not surprising, as compromised cellular signaling is a hallmark of cancer cells (Sever R. et al 2015). In addition, a total of 231 mutations fall in the protein domains of 137 tumor proteins. Overall, a large number of proteins are modified at multiple residues in this data set; also, many proteins suffer from the modified residue mutations. If further explored, this dataset can aid in distinguishing and interpreting a specific type of tumor with a specific repertoire of PTM-mutational load.

In the second case, I performed Pyntheon analyses on proteins that are linked to plant *Arabidopsis thaliana* disease/stress resistance (Haung et al 2010). The use of a Pyntheon command (B) showed that proteins that are associated to the plant resistance are modified at different residues and that these modifications have a potential role to play in the resistance (see supplementary data).

### $ python3 pyntheon.py –f file.txt –s at -p TRUE…… (B)

These examples show the potential of Pyntheon in evaluating the involvement of protein functional features in a phenotype, in addition to the robustness of it in performing such analyses for a large number of species. For an explanation of the commands used, the data, and the presented in these examples, see the Code-Availability.

## 3 Conclusion

The development of such a comprehensive tool will generate interest in analysis of mutational overload impacting the PTMs present in a large number of species. The ease of use for the analyses of large experimental datasets will accelerate the understanding and interpretation of PTM functions-dependent biological systems. I believe, Pyntheon will help the larger scientific community study PTMs in different species.

In future development, I am aiming to implement Pyntheon in R and in an HTML-based user interface, to further facilitate users.

